# Rapid Sensing of Volumetric Neural Activity through Adaptive Remote Focusing

**DOI:** 10.1101/125070

**Authors:** Mantas Žurauskas, Oliver Barnstedt, Maria Frade-Rodriguez, Scott Waddell, Martin J. Booth

## Abstract

The ability to record neural activity in the brain of a living organism at cellular resolution is of great importance for defining the neural circuit mechanisms that direct behavior. Here we present an adaptive two-photon microscope optimized for extraction of neural signals over volumes in intact *Drosophila* brains, even in the presence of specimen motion. High speed volume imaging was made possible through reduction of spatial resolution while maintaining the light collection efficiency of a high resolution, high numerical aperture microscope. This enabled simultaneous recording of odor-evoked calcium transients in a defined volume of mushroom body Kenyon cell bodies in a live fruit fly.

---

Investigating functional connectivity between neurons in the brain is the key to understanding many fundamental questions in neuroscience. Recording neural activity with genetically encoded calcium indicators such as GCaMP^1^ permits non-invasive observation of multiple neural responses, if combined with a suitable microscopic method. Yet, imaging activity quickly over large volumes inside a live brain still remains one of the greatest challenges in neuroscience microscopy^2,3^. Images from widefield single photon fluorescence microscopes can encompass a large number of neurons, but the cells of interest do not necessarily lie in one image plane. Moreover, the volumetric nature of neural tissue means that out-of-focus background often swamps useable signals. Scanning two-photon microscopy is commonly favored for imaging in scattering neural tissues as it provides high contrast, deep penetration, increased resilience to scattering and low phototoxicity to the sample. However, to compile a volume multiple planar images need to be acquired. Collecting data in this way is slow because the specimen or objective has to be moved. Imaging-induced and natural motion of the specimen are problematic because the plane of interest can be easily lost from view – our own investigations have revealed frequent translations of several micrometres when imaging neurons in the *Drosophila* brain. Single objective scanning light sheet methods could be a solution, although resolution and collection efficiency are not optimal due to the limited numerical aperture^4^. Light field microscopy can image large volumes simultaneously, but it relies on sparse labeling^5^. Two-photon fluorescence methods present the most promising avenue for fast functional imaging of densely labeled specimens. Here we have taken a new approach to fast two-photon imaging – Adaptive Optics based Volumetric Activity Sensing Two-photon (AO-VAST) microscopy – that is optimized for extraction of information from dense neural tissue.

Volumetric imaging rates can be increased by avoiding the need to move the specimen or objective. To this end, various optical axial scanning techniques have been employed. The remote-focusing principle, which is common to all of these high-speed volumetric techniques, involves shifting the imaged plane by application of a defocus phase at the pupil of the optical system. The fastest available methods rely on acousto-optical scanning for 3D random access of multiple pre-defined points within the sample^6,7^. These methods are optimized for sampling of multiple spatially dispersed points, but are not practical for full volume imaging. Furthermore, the focusing is not ideal, as the quadratic phase functions introduced by the acousto-optical scanners do not perfectly match the phase required for spherical-aberration-free focusing. Systems using matched lenses and small mirror based refocusing have been used to scan in 3D at line rates of several kHz with aberration-free focusing, but the speed is still limited for complete volumes^8^. Other techniques include electrically tunable liquid lenses^9^, which are slow and do not provide aberration free focusing, or ultrasound driven lenses^10^, which again suffer from focusing aberrations and are limited to fast sinusoidal scanning. AO-VAST microscopy provides a combination of capabilities that cannot be achieved with these existing technologies.

Measuring brain activity with high temporal resolution requires a new design philosophy to exploit different attributes of available technologies. Foremost, prioritizing the extraction of information about cell activity over the resolution of structural details leads to very different specifications for the imaging system. AO-VAST combines several technologies to enable this rapid 3D sensing of neural activity (Fig 1a). Line scanning two-photon microscopy^11^ provides higher scanning speed than a point scanning approach, when employing standard off-the-shelf galvanometric mirror systems. This is combined with non-descanned detection on a sCMOS camera. Adaptive optics (AO) based remote focusing is enabled by a deformable mirror (DM), which is positioned conjugate to the objective pupil plane. The DM serves two purposes: sample-induced aberration correction and shifting simultaneously the focal position of the excitation and imaging paths^12^. The line scanning strategy combined with AO axial scanning, allows speeds of up to 1000 planes, or 100 volumes per second when using strongly fluorescent samples (Fig. 1f).

**Figure 1.**
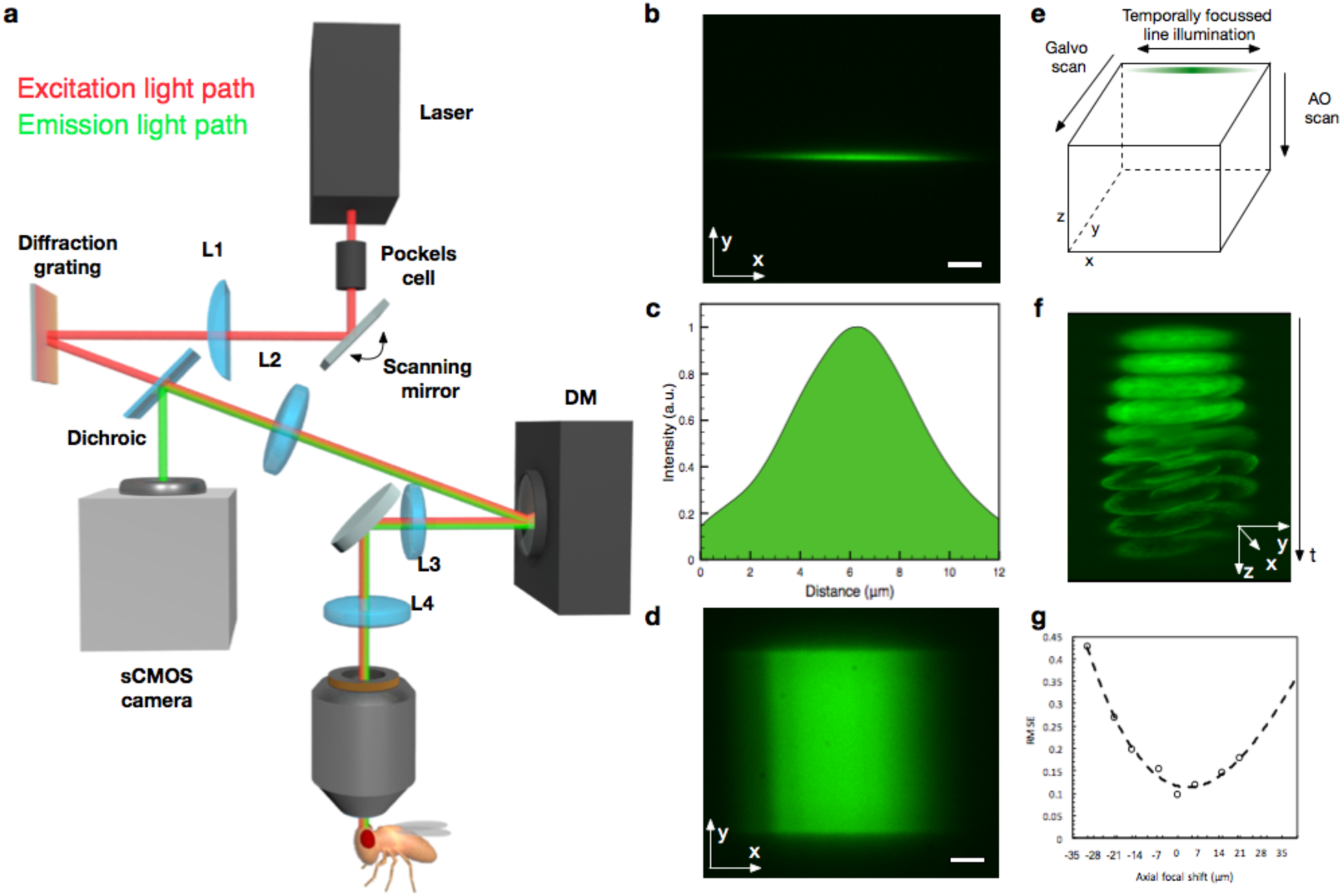
**(a)** Simplified schematic diagram of the AO-VAST microscope configured for *in vivo* imaging. **(b)** Illumination line exciting two-photon fluorescence in a fluorescent polymer slide indicating the shape of line illumination profile. **(c)** Axial profile of the excitation PSF obtained by recording fluorescence intensity when a thin layer of fluorescent dye is scanned through object plane **(d)** A single imaging plane obtained by laterally scanning an excitation line across a fluorescent plastic slide. **(e)** Volumetric imaging strategy: each optical slice is formed by scanning a line along the image plane; scanning along the optical axis is enabled by a deformable mirror. **(f)** Expanded 3D image of a pollen grain: the whole volume consisting of ten z-slices, each of thickness ∼5 μm, was captured in 10 ms. **(g)** Wavefront root mean square error (RMSE) when focusing at different axial positions. Data points are fitted with a 2^nd^ order polynomial. In **b** and **d** the scale bar corresponds to 10 micrometers.

Temporal focusing has been used previously in the context of two-photon imaging to decouple axial and lateral confinement of excitation^13^. We took advantage of this by using temporal line focusing to match the excitation profile in the z direction to the typical size of Kenyon Cell bodies (Fig. 1c). Kenyon Cell somata are relatively large (up to 5 μm) compared to the resolution of a standard two-photon microscope (about 0.5 μm) and are usually distributed over a volume. Furthermore temporal focusing has been shown to provide increased robustness of the excitation beam against the effects of scattering^14^. Additionally, depending on imaging requirements, the scanning speed can be traded for a reduction of the illumination beam average power, possibly leading to reduced phototoxicity^15^ to the sample.

A state of the art deformable mirror (DM) can control the wavefront at rates of up to 20 kHz^16^. This speed is more than adequate when measuring even the fastest commonly used calcium or voltage indicators^1^. The DM is also able to produce more complex shapes than the quadratic phase function that is offered by many simpler focusing methods. This is necessary, as refocusing in high numerical aperture systems requires a high order polynomial – not quadratic – phase function for aberration free operation. Furthermore, the DM can be used to compensate sample and system induced aberrations leading to increased signal to noise ratio (SNR) that permits shorter exposure times. DM based systems have the advantage of compactness and have a potential to exhibit lower optical loss than acousto-optical^6,7^ or lens based^8^ remote focusing systems with similar functionality. Realistic labelling densities mean that SNR is the limiting factor for fast imaging. However, SNR can be increased if the spatial resolution of the microscope is intentionally reduced so that, in effect, an image voxel measures the signal integrated over the region of interest, such as a cell body. Fast monitoring of continuous volumes can therefore be achieved by increasing the voxel size to approach the physical dimensions of a cell body, whilst maintaining the fluorescence collection efficiency of a high NA objective lens. The resulting optical sections are thicker than standard two-photon microscopes, thus fewer images – and less time – are needed to cover the whole volume. The method also avoids the problematic loss of data due to motion of the live specimens – since movements are contained within the imaged volume, they can be dealt with through post-processing. Note that an equivalent effect cannot be achieved by using an objective lens with lower numerical aperture – while the axial resolution can be reduced this way, the fluorescence detection efficiency drops dramatically with the lower aperture angle.

## Results

To characterize the deviations from the ideal optical system performance^17^ and to measure *in situ* the phase maps required to shift the focal plane in an aberration-free manner, we have used custom wavefront measurement based on solution of the transport of intensity equation^18^. A lookup table obtained during the mirror training **(Supplementary Fig. 3)** was then used to shift the focal plane (Fig. 1d) to the required axial position at up to kHz rates. In a double pass configuration, a single DM permits refocusing for both excitation and fluorescence emission paths. For wavefront sensing and mirror training methods, please refer to **Methods section**.

Combined, these technologies can deliver fast and robust neural activity sensing *in vivo*. We imaged activity in somata of the Kenyon cells in the mushroom body in awake *Drosophila* using the genetically encoded calcium indicator GCaMP6f (Fig 2). Rapid monitoring of Kenyon cell body activity is difficult with current single plane imaging techniques, as the cell bodies are distributed throughout the volume and they respond simultaneously in different axial positions.

**Figure 2.**
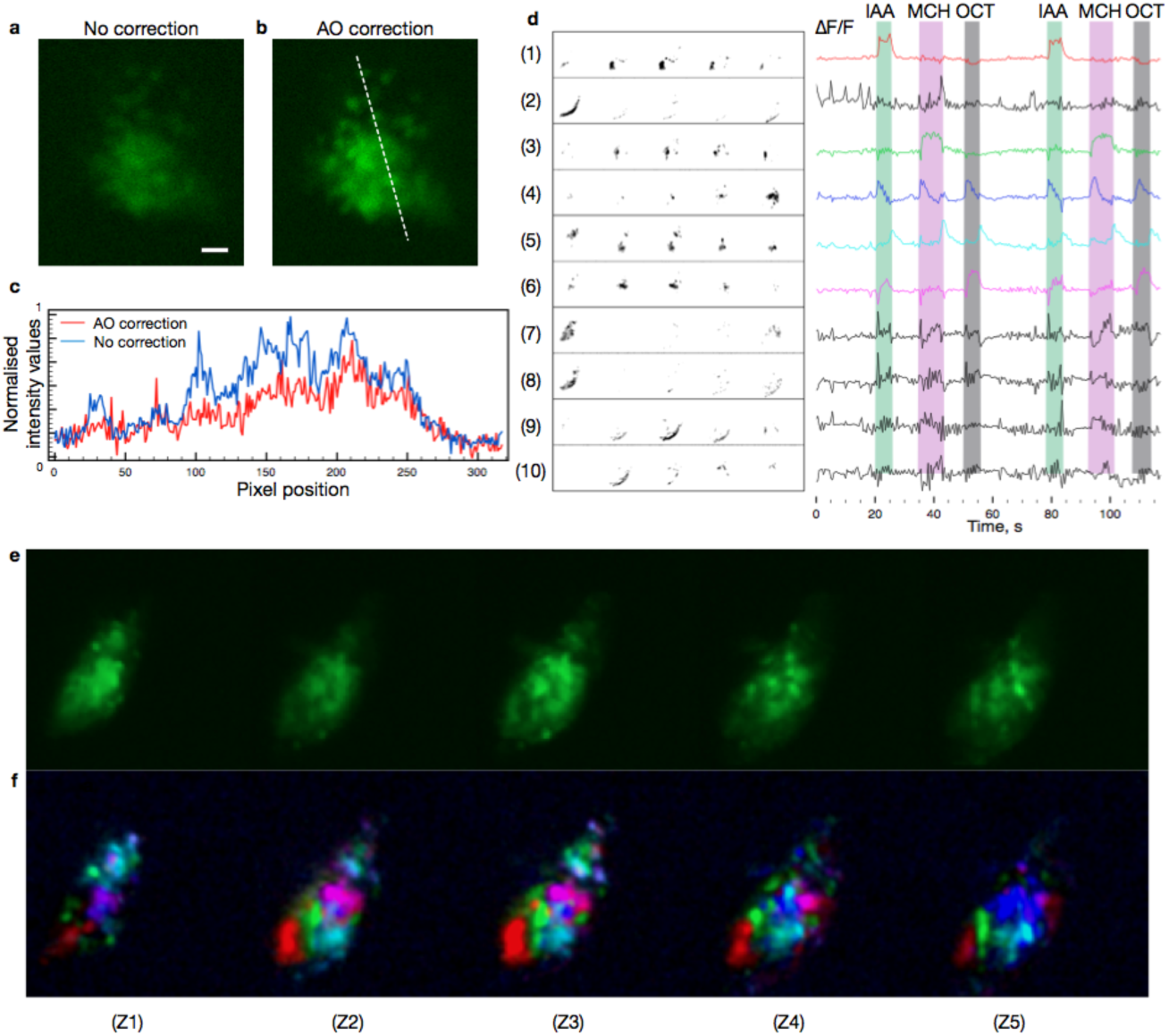
Imaging odor responses in a live fruit fly brain expressing GCaMP6f in mushroom body Kenyon Cells. **(a)** Image of Kenyon Cell bodies with system flat correction. Scale bar corresponds to 10 micrometers. **(b)** AO correction of sample induced aberrations leads to sharper images and increased image brightness. **(c)** Comparison of raw images acquired with and without correction of sample induced aberrations. AO correction leads to increased signal to noise ratio, permitting faster imaging. **(d)** Independent components extracted from a recorded video (**Supplementary Video 1**) and ΔF/F responses corresponding to each independent component. Green, purple and grey color bars represent the time periods where the fly was exposed to IAA, MCH and OCT odors. **(e)** Averaged recording of Kenyon Cells expressing GCamp6f representing cumulative fluorescence throughout full odor delivery sequence. Each image going from left to right corresponds to a 50x80 μm consecutive z-slice separated by 5 μm. **(f)** A composite image where different colors correspond to spatial filters 1,3,4,5,6 in **(d)**.

As a demonstration of capability, we imaged a volume of 50x80x20 μm consisting of a set of 5 axial slices (each about 5 μm thick) separated by 5 μm, being imaged at a rate of 2 volumes per second. The instrument design meant that no gaps existed between adjacent sections. Excitation power was set to 100 mW average power spread across the excitation line (Fig. 1c). During imaging, the objective and the sample were kept in fixed positions. Even though there were obvious movements during acquisition, no pre-processing for movement correction was performed to demonstrate the robustness of the method. The approach could be extended to more extreme movements by pre-processing to realign the images in three dimensions.

We also extended automated cellular signals analysis methods^19^ to work with volumetric data to identify the groups of cells that respond together during the stimulation. The resulting ΔF/F traces (Fig. 2) show that our fast sensing method can reproducibly capture the responses of cell groups that respond to different odor stimuli. Furthermore, initial results indicate that our method may be used for long periods without causing significant phototoxicity to the fly. A video **(Supplementary Video 2**) shows a fly responding to odor stimuli while Kenyon Cell bodies are imaged in different locations with laser power varying from 70 mW to 140 mW for over 25 minutes. We have noticed that, depending on the quality of sample preparation and the power of laser excitation, the imaging time can be easily extended up to an hour. Defining the limits will require more experiments.

The AO-VAST sensing approach enables monitoring of neural activity throughout continuous volumes. The combination of temporal focus line scanning two-photon excitation and AO based remote focusing permits adjustable reduction of spatial sampling over the volume in order to increase temporal resolution. As a result of these resolution compromises, one can extract enough information to determine volumetric activity patterns at the rates of up to 100Hz that are ultimately limited by DM speed and sample brightness. These rates are not achievable with standard point scanning two-photon imaging modalities. Furthermore, the DM used for fast remote focusing, can be simultaneously employed to cancel sample induced aberrations and further increase SNR to boost practical imaging rates. Lastly, the compact footprint of the system allows it to be easily added to commercial two-photon microscopes (**Supplementary Fig. 1b**).

## Methods

### *In vivo* imaging of odor responses in a *Drosophila* brain

The optical set-up for the AO based volumetric activity sensing two-photon (AO VAST) microscope (**Supplementary Fig. 1**) was built in-house and mounted on top of a Scientifica two-photon microscope (**Supplementary Fig. 2**). The optical power of a femtosecond excitation laser (Chameleon Ultra II, Coherent Inc.) was controlled by adjusting the voltage applied to a Pockels cell (350-80, ConOptics). The collimated laser beam was launched on a scanning mirror (GVSM001, Thorlabs) positioned at the pupil plane of the optical system, the beam was then focused with a cylindrical lens (f=100mm) to form a line on a grating (1200 lp/mm ruled grating, Edmund Optics), which was used to enable spatio-temporal focusing. For optimal diffraction efficiency, the laser polarization was aligned to the grating by manually adjusting the rotation of a half wave plate HWP. The light from the grating was then relayed on to a deformable mirror (DM-69, Alpao) positioned at the pupil plane of the system and re-imaged on to the objective lens (40× water immersion LUMPLFLN 40XW, Olympus).

The fluorescence light propagating back through the same path was separated from illumination beam just after the lens L2 using a dichroic beam splitter (FF665Di02, Semrock), and de-magnified with a matched achromatic lens pair (1:3.3, Thorlabs). The image was then captured with an sCMOS camera (Zyla 4.2P, Andor).

The system permits convenient switching between arc-lamp illumination (Xcite 120PCQ, Lumen Dynamics) for locating the area of interest and two-photon illumination for fast 3D imaging by rotating the filter wheel which is built in the Scientifica two-photon microscope stand.

### Deformable mirror training for remote focusing

To enable remote focusing, a lookup table containing phase maps corresponding to different focal positions was built (**Supplementary Fig. 4**). For each focal position, the optical system was characterized by measuring the wavefront of the light propagating through the system (**Supplementary Fig. 3)** with a flat mirror positioned in the sample plane. During the characterization, the system was switched to wavefront sensing configuration (**Supplementary Fig. 1**).

### Odor delivery

For the odor response imaging, three different odors were delivered on a clean air carrier stream using a custom-designed system^20^. 5 seconds of IAA (isoamyl acetate) were followed by 12 seconds of air, then 8 seconds of MCH (methylcyclohexanol), then 10 seconds of air, and finally 5 seconds of OCT (octanol).

### Fly preparation for imaging

For live imaging we used LexAop-GCaMP6f; 247-LexA::VP16 flies^20,21^. Three-to eight-day-old flies were briefly anesthetized on ice and mounted in a custom chamber. The head capsule was opened under room temperature carbogenated (95% O2,5%CO2) buffer solution (103 mM NaCl, 3 mM KCl, 5 mM N-Tris, 10 mM trehalose, 10 mM glucose, 7 mM sucrose, 26 mM NaHCO3,1 mM NaH2PO4, 1.5 mM CaCl2, 4 mM MgCl2, osmolarity 275 mOsm [pH 7.3]). The legs and proboscis were immobilized with wax.

### Data format and analysis

Raw image data were stored as TIFF images for each plane in the series of imaged volumes. The saved data were processed with an automated analysis algorithm^19^ adapted for processing 3D data. For the processing, the frames corresponding to all depths depth in a single volume were stitched together side-by-side. This allowed a single volume to be considered as a single frame. The spatial modes that correspond groups of cells that respond together were identified by performing automated sell sorting in four steps: Principal component analysis, spatio-temporal independent component analysis, image segmentation and deconvolution. Filters based upon the spatial modes were then used to extract the time traces that represent the dynamics of the cell activity.

ImageJ was used to crop the images, and to adjust brightness and contrast for display purposes, and for 3D rendering.

### Funding

Medical Research Council (MR/K01577X/1); S.W. is funded by a Wellcome Trust Principal Research Fellowship in the Basic Biomedical Sciences and the Bettencourt-Schueller Foundation.

